# Inducing Human Retinal Pigment Epithelium-like Cells from Somatic Tissue

**DOI:** 10.1101/2020.07.27.215103

**Authors:** Ivo Ngundu Woogeng, Imad Abugessaisa, Akihiro Tachibana, Yoshiki Sahara, Chung-Chau Hon, Akira Hasegawa, Bogumil Kaczkowski, Noriko Sakai, Mitsuhiro Nishida, Haiming Hu, Hashimita Sanyal, Junki Sho, Takeya Kasukawa, Minoru Takasato, Piero Carninci, Akiko Maeda, Michiko Mandai, Erik Arner, Masayo Takahashi, Cody Kime

**Affiliations:** RIKEN Center for Biosystems Dynamics Research, Kobe 650-0047, Japan; RIKEN Center for Integrative Medical Sciences, Yokohama 230-0045, Japan; Laboratory of Molecular Cell Biology and Development, Department of Animal Development and Physiology, Graduate School of Biostudies, Kyoto University, Kyoto 606-8501, Japan; Department of Renal and Cardiovascular Research, New Drug Research Division, Otsuka Pharmaceutical Co. Ltd., Tokushima 771-0192, Japan

**Keywords:** cell biology, cell plasticity, reprogramming, regenerative medicine, retina

## Abstract

Regenerative medicine relies on basic research to find safe and useful outcomes that are only practical when cost-effective. The human eyeball requires the retinal pigment epithelium (RPE) for support and maintenance that interfaces the neural retina and the choroid at large. Nearly 200 million people suffer from age-related macular degeneration (AMD), a blinding multifactor genetic disease among other retinal pathologies related to RPE degradation. Recently, autologous pluripotent stem cell-derived RPE cells were prohibitively expensive due to *time,* therefore we developed a new simplified cell reprogramming system. We stably induced RPE-like cells (iRPE) from human fibroblasts by conditional overexpression of broad plasticity and lineage-specific pioneering transcription factors. iRPE cells showed features of modern RPE benchmarks and significant *in-vivo* integration in transplanted chimeric hosts. Herein, we detail the iRPE system with comprehensive modern single-cell RNA (scRNA) sequencing profiling to interpret and characterize its best cells. We anticipate that our system may enable robust retinal cell induction for regenerative medicine research and affordable autologous human RPE tissue for cell therapy.

## INTRODUCTION

The retinal pigment epithelium (RPE) is a monolayer of cuboidal cells developed between the photoreceptors and choroid of the eye. The RPE is critical for the development, maintenance, and function of photoreceptors, and mutations in key RPE genes may cause degenerative retinal disorders such as retinitis pigmentosa (RP) with a prevalence of 1 in 4000 people as the most common inherited retinal dystrophy (Esumi et al., 2004; Verbakel et al., 2018). Supplementation of RPE cells can recover from RPE dysfunction in animal models, suggesting a potential solution for RP (Haruta et al., 2004; Maeda et al., 2013). RPE degeneration onset is also associated with preceding age-related macular degeneration (AMD), the leading cause of irreversible blindness in western countries (Esumi et al., 2004; Klein et al., 1992; Smith et al., 2001). In AMD, the RPE cells are invariably lost, therefore intervening AMD treatments to prevent blindness may require cell transplant therapy (Mandai et al., 2017).

Recent advances in regenerative medicine have motivated new strategies to develop pluripotent stem cell-derived RPE from human embryonic stem (ES) cells and autologous induced pluripotent stem cells (iPSC) (Haruta et al., 2004; Mandai et al., 2017). RPE differentiation from pluripotent stem cells was pioneered against great technological and political barriers that were overcome with demonstrably safe and functional iPSC.RPE for patient transplant (Mandai et al., 2017). Still, uncertainties about iPSC potency and genome stability remain, and such novel regenerative medicine required labor and financing that are impossible to budget in modern health systems. Therefore, simplifying the induction of autologous RPE cells with a more direct approach may be necessary.

Lately, ‘direct reprogramming’ systems that convert between somatic cell states (Najm et al., 2013; Pang et al., 2011; Zhang et al., 2014) have emerged, inspired by the transcription factor (TF) synergy famously uncovered by Kazu Takahashi and Shinya Yamanaka with iPSCs (Takahashi et al., 2007). Such systems posit that core TF sets should ‘directly’ convert between somatic cell identities (D’Alessio et al., 2015; Rackham et al., 2016). Conceptually, ‘direct reprogramming’ ignores epigenetic plasticity, or assumes it, with simplistic design for somatic to somatic cell state conversion. In practice, such systems usually rely on Yamanaka-factor (OCT4, SOX2, KLF4, MYC) co-induction, or transit cell progenitor and intermediate plastic states yet bound by *time* and a *surviving identity.* Unlike somatic cell identities, induced pluripotency enables increasing plasticity via pioneering TF driven epigenetically self-recursive state reinforcement, termed ‘maturation’, that was obviated much later (Iwafuchi-Doi and Zaret, 2014; Samavarchi-Tehrani et al., 2010; Soufi and Zaret, 2013; Soufi et al., 2012). Indeed, once minimally established, *in-vitro* pluripotency self-iterates and stabilizes its plastic pluripotent identity. To systematically induce RPE (iRPE) from fibroblasts, TFs with pioneering activity, direct roles in plasticity, and the developmental differentiation and specification of RPE may be required (Soufi et al., 2015).

To reduce the costly *time* for autologous cell production, we looked to engage epigenetic plasticity at the same time as an induced RPE state using the aforementioned criteria and found that four TFs, enhanced by CRX and small molecules, could convert human fibroblasts to bulk cultures containing RPE-like cells with characteristic function, expression, cell identity, and integration in chimeric subretinal transplants. Together, this iRPE platform and scRNA datasets might be used to develop affordable autologous biomedical-grade regenerative RPE cell therapies.

## RESULTS

### Few Exogenes Are Necessary to Reprogram Human Somatic Cells to RPE-like Fate

Inducing autologous iPSC-derived RPE (iPSC.RPE) has aided cell therapy studies (Mandai et al., 2017) with a multiplier for costs based on *time*. For practicality, we set out to induce human somatic cells much more quickly to RPE cells (Figure 1A). We looked to a previous ‘direct reprogramming’ study (Zhang et al., 2014), but such factors failed in our hands (data not shown). However, we found that a conditionally expressed combination of RPE cell-specific (MITF, CRX, OTX2), lineage-specific (CRX, OTX2), and pluripotency/plasticity/regenerative genes (OTX2, LIN28, MYCL) could rapidly induce human foreskin fibroblasts (Fib) to RPE-like cells with tacit features of RPE summarized in Figure 1. We employed lentiviral transduction to introduce a molecular toolset containing a common minimized BEST1 (VMD2) (Esumi et al., 2004; Masuda and Esumi, 2010; Zhang et al., 2014) synthetic reporter construct to drive EGFP expression (*BEST1::*EGFP) and a constitutive polycistronic sequence with dox-inducible rTTA and Puromycin resistance (Figure 1B). Fib were transduced with the toolset, selected briefly with Puromycin, expanded, and then transduced with dox-inducible TetRE transgenes (Figure S1A). Importantly, the *BEST1::*EGFP reporter responded to iPSC.RPE cell maturity and density, with similar expression in our RPE-like cells. We sub-cultured picked colonies of such induced iRPE cells and validated RPE65 protein expression among the EGFP+ cells (Figure 1C). We termed these induced RPE-like cells iRPE.

**Figure 1:**
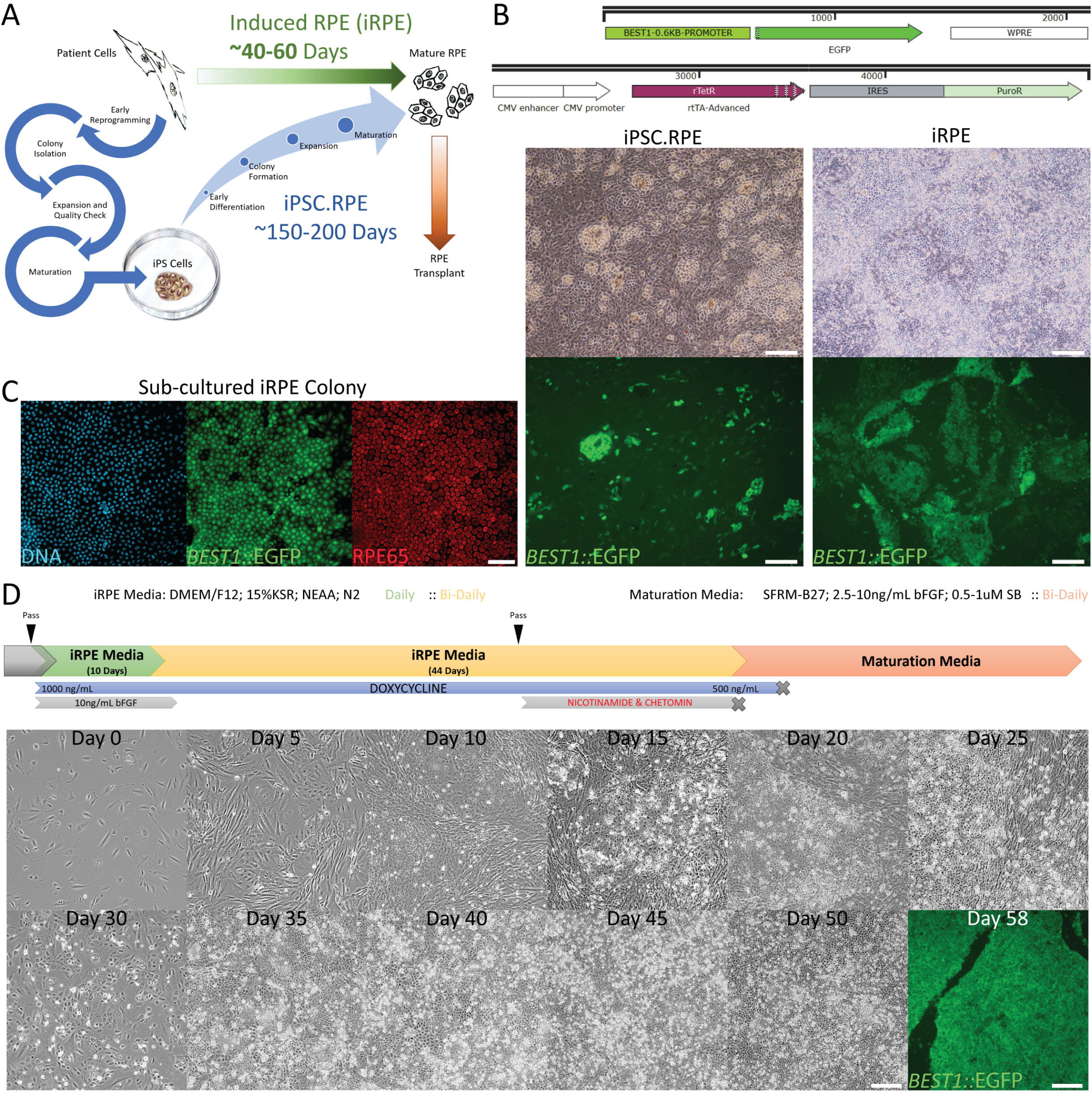
iRPE System Overview. **A)** A schematic representing the iRPE system objectives and estimated time frames when comparing autologous iRPE to autologous iPSC.RPE. **B)** The *BEST1::*EGFP synthetic promoter reporter construct and constitutive (CMV) driven conditional dox-inducible system (rTetR/rtTA) with Puromycin Resistance (PuroR). Construct is integrated and expressed in iPSC.RPE (left) and iRPE (right). *Scale bars = 200μm.* **C)** A sub-cultured iRPE colony expressing *BEST1::EGFP* stained for DNA (light-blue; Hoechst 33342) and RPE65 (red). *Scale bar = 100μm.* **D)** A schematic (upper) and pictorial (lower) representation of iRPE reprogramming with basal media compositions and supplementations, timing of molecules and conditional reprogramming (doxycycline). *Scale bars = 100μm.*

Individually sub-cultured iRPE colonies did not proliferate or expand much past 0.64 cm^2^ (Figure S1B), and thus we usually bulk passaged our full 6W well 1:2 on approximately Day 28-30 (Figure 1D). With this system, we generally observed distinct morphological change and mesenchymal to epithelial transition (MET) between Days 5 and 10 and specific cobblestone RPE-like morphology with variable activation of *BEST1::*EGFP between Days 12 and 25 and variable stability after removal of doxycycline (Figure 1D). Taken together, these observations reinforced the notion that our iRPE system may transition through important MET mediated cell identity reprogramming with genome stability selectivity (Kareta et al., 2015; Li et al., 2010; Marión et al., 2009; Samavarchi-Tehrani et al., 2010), and then acquire important features and reporters of RPE cell identity (Maruotti et al., 2015; Masuda and Esumi, 2010; Zhang et al., 2014).

### Orthodenticle Genes are Powerful Effectors of iRPE Reprogramming

Previous reports had resolved or hypothesized (Rackham et al., 2016) iRPE system factors (Figure 2A), but did not turn off reprogramming factor expression (Zhang et al., 2014), or observed a return to Fib identity when doing so (D’Alessio et al., 2015). Both reports that demonstrated iRPE cell output utilized Orthodenticle homeobox 2 (OTX2), a powerful gene expressed from arrangement and induction of primed pluripotency through development to the eye, and then the RPE (Buecker et al., 2014; Esumi et al., 2009; Hever et al., 2006; Thomson and Yu, 2012).

**Figure 2:**
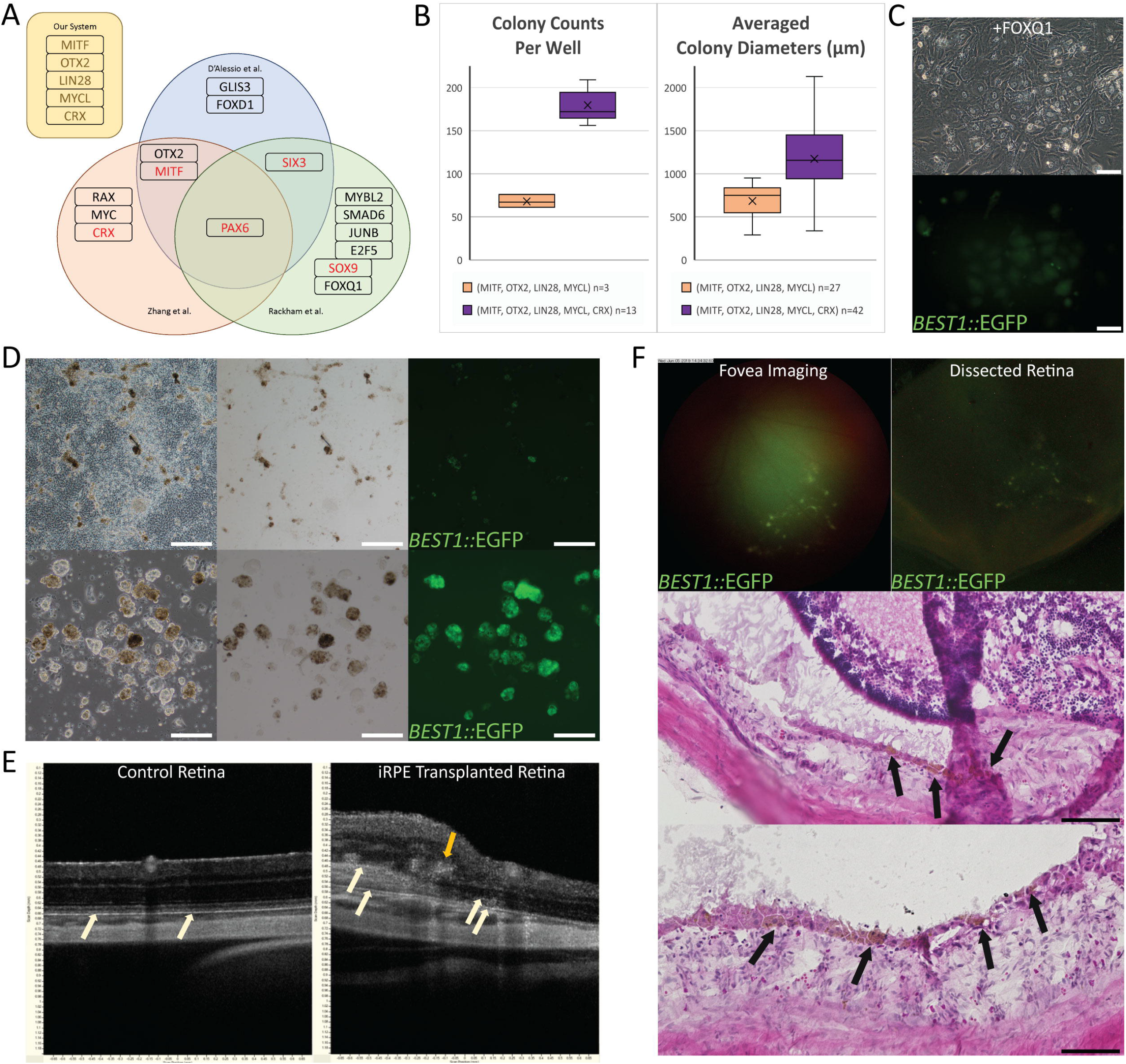
iRPE Systems, TF Testing, and Preliminary Subretinal Transplantation. **A)** A Venn-diagram of the validated TFs from our iRPE system and each previous iRPE study (D’Alessio et al., 2015; Rackham et al., 2016; Zhang et al., 2014). **B)** iRPE reprogramming +/− CRX, with colonies counted per 6W on Day 9 (left) and average colony diameters measured on Day 13 (right). **C)** Day 9 iRPE system +FOXQ1 colony expressing *BEST1::*EGFP. *Scale bars = 100μm.* **D)** Pre (upper) and Post (lower) iRPE cluster purification culture images to show morphology, pigmentation, and *BEST1::*EGFP expression. *Upper scale bars = 500μm, Lower scale bars = 200μm.* **E)** Optical Coherence Tomography scans of untreated albino rat retina (left) and iRPE transplanted albino rat retina (right). Host and transplanted RPE/RPE-like layers are indicated (white arrows) along with potential retinal transplant rosette formations (yellow arrow). **F)** Live fovea fluorescent imaging for *BEST1::*EGFP (upper left), with the same explanted dissected retina fluorescent imaging for *BEST1::EGFP* (left), and that retina’s cryosections with H&E staining to reveal pigmented human iRPE cells (black arrows) in RPE layers and subretinal space, interfacing with host photoreceptor outer segments. *Scale bars = 100μm.*

A notorious retinal development TF, Orthodenticle homeobox 3, is commonly called Cone-Rod homeobox protein (CRX) (Esumi et al., 2009). In early tests, we isolated iRPE colonies for expansion and performed genomic DNA PCR for our reprogramming transgenes and found that MITF was not common yet the Orthodenticle homeobox TFs CRX and OTX2 were detected in all clones (Figure S1C). We found that excluding CRX resulted in significant reductions in iRPE colony counts per well and smaller colony diameters (Figure 1B). We therefore maintained CRX and OTX2 in our core iRPE system experiments.

Among all previous iRPE reports (Figure 2A), and many other ‘direct reprogramming’ systems, the pioneering TF PAX6 (Soufi et al., 2015) was common, and we anticipated an improvement to our system. We added dox-inducible PAX6 to our reprogramming set (MITF, OTX2, LIN28, MYCL, CRX) and observed a broad increase in cell morphology change by Day 3 followed by a striking cell death event and total ablation of iRPE colony forming cells by Day 6, leaving no visible colonies for an extended period thereafter (data not shown). Alternative iRPE reprogramming factors FOXQ1 and SOX9 were also added to our system. However, both factors caused premature activation of the *BEST1::*EGFP reporter, rendering its RPE maturation/identity features useless (Figure 2C). Furthermore, FOXQ1+ reprogramming induced small EGFP+ colonies with little to no visible proliferation by Day 9 (Figure 2C). For these reasons, we did not continue to apply PAX6, FOXQ1, or SOX9 from the beginning of iRPE reprogramming.

### Transplanted iRPE Cells Integrate and Pigment in Albino Rat Retinas

In preliminary tests, *BEST1::*EGFP+ reporting iRPE cells in experiments were collected in floating pigmenting balls for purification similar to our previous iPSC.RPE production method (Kuroda et al., 2012). We pooled several floating pigmenting iRPE clusters to a well and sub-cultured for brief expansion (Figure 2D). We prepared immune-compromised Albino rats with subretinal transplantation of iRPE cells. Within 2-3 months we used Optical Coherence Tomography (OCT) scans and usually found several affected areas with possible bulks of cells between the host RPE and Photoreceptors. Notably, some cells in the bulked areas had structural and light characteristics that resembled the RPE layers. We also saw signs of xenografted cells in the photoreceptor layer, implicating rosettes often seen in RPE xenograft experiments (Figure 2E, Figure S2A).

We observed a weak trace of EGFP+ cells during fluorescent live fovea imaging, and fluorescent imaging of all transplanted dissected retinas (Figure 2F). Cryosections of those retinas, prepared with hematoxylin & eosin (H&E) staining, showed that many pigmented cells were cobblestone-morphology and found interfacing with photoreceptor outer segments. These cells were sometimes fully integrated into the RPE layer at various positions proximal to the injection site (Figure 2F).

### Nicotinamide and Chetomin Improves iRPE Cell Reprogramming

Previous reports showed that Nicotinamide (NIC) and Chetomin (CTM) treatments may improve pluripotent stem cell-derived RPEs, such as iPSC.RPE (Maruotti et al., 2015; Williams et al., 2012). We performed bulk passage of iRPE cells to two wells and treated one well +CTM mid-reprogramming with the timing shown in Figure 1D. iRPE+CTM appeared to reduce *BEST1::*EGFP maturation reporter expression and cell pigmentation among significant cell debris/death while the surviving cobblestone cell layer became morphologically more homogenous than control cells (Figure 3A). Next, we prepared trans-well cultures with human primary RPE (hRPE), iRPE, and iRPE+CTM. We examined apical and basal PEDF and VEGF concentrations by ELISA, along with TER measurements, across a 4-week period (Figure S3A). Generally, iRPE and iRPE+CTM did secrete PEDF and VEGF with apical/basal secretion trends like the hRPE, although weaker. TER measurements also showed a lower initial TER, and 4-week increase, when compared to hRPE. Interestingly, iRPE+CTM samples were improved over iRPE alone (Figure S3A).

**Figure 3:**
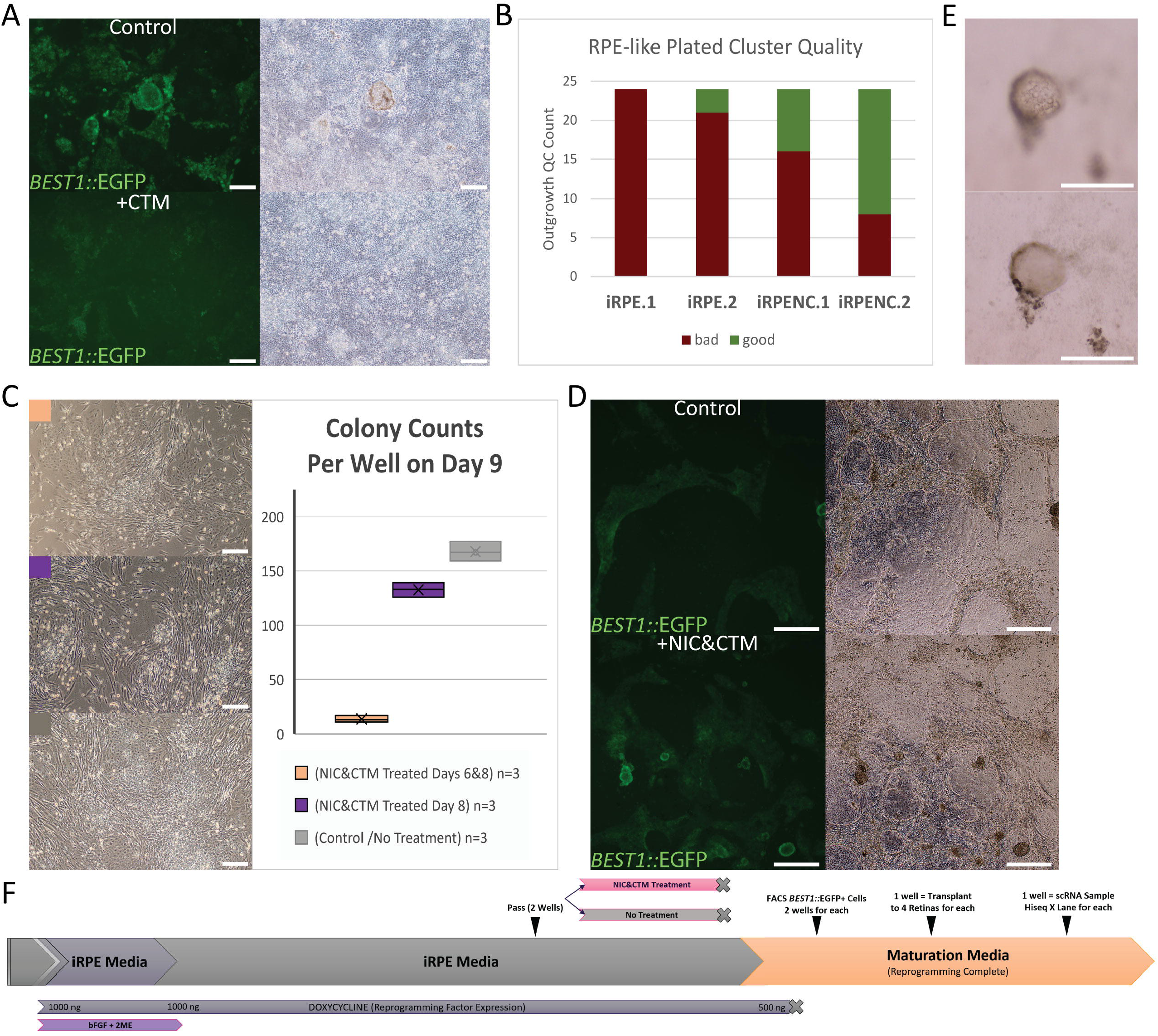
iRPE Reprogramming Cells Treated with Nicotinamide and Chetomin. **A)** A split iRPE culture +/− CTM treatment, during treatment, with *BEST1::*EGFP expression (left) and brightfield imaging (right). *Scale bars = 200μm.* **B)** Subjective qualification (bad/good) of 24 individually plated iRPE cluster outgrowths from 2 iRPE and 2 iRPENC cultures. **C)** Effects of NIC&CTM treatments on early iRPE reprogramming cultures shown with brightfield microscopy (left) and iRPE colony counts on Day 9 (right). *Scale bars = 200μm.* **D)** A split iRPE culture that had previous +/− NIC&CTM treatments, prior to retinal transplant, expressing *BEST1::*EGFP among variable quality cells visible by brightfield (right). *Scale bars = 500μm.* **E)** A representative bulging RPE-like bleb of iRPE cells imaged at two z positions (upper/lower) with obviated pigmentation and cell nuclei (upper). *Scale bars = 200μm.* **F)** A schematic overview of how the iRPE and iRPENC cells were prepared for scRNA analysis and albino rat subretinal transplant.

To examine the effects of NIC and CTM to the iRPE system, we performed a larger bulk passage of the same cells to several wells treated with NIC, CTM, or NIC&CTM (Figure S3B). Interestingly, NIC alone had increased *BEST1::*EGFP expression over the control samples during treatment and afterward such cells had increased cell pigmentation and bulging/blebbing as cells matured. CTM alone followed the previously observed trend, decreasing *BEST1::*EGFP expression during treatment, and had mild reduction in cell pigmentation as cells matured. Excitingly, the combination of NIC&CTM appeared to gain the benefits of each individual treatment with no notable negative effects. We therefore termed iRPE +NIC&CTM as iRPENC and generally use that treatment as standard (Figure 1D).

During RPE purification, individually plated pigmented clusters can reveal a subjective basic quality based on outgrowth morphology, as done with iPSC.RPE (Kuroda et al., 2012). We performed parallel experiments of iRPE and iRPENC originating from the same bulk passage, but post-maturation and doxycycline-removal. Purified pigmented floating clusters were then plated to individual wells, and outgrowths were assessed as ‘good’ or ‘bad’. We observed that the iRPENC cultures produced significantly more ‘good’ outgrowth cultures (Figure 3B).

We noted an interesting selective cell death from CTM treatments in iRPE reprogramming that was not described in the previous iPSC.RPE differentiation study (Maruotti et al., 2015). Indeed, IRPE reprogramming is uniquely from fibroblasts, and involves a bulk passage such that non-iRPE cells are outcompeted but may remain. We tested NIC&CTM treatments in early iRPE reprogramming and found that fibroblasts and early iRPE colony formation were drastically affected. Interestingly, if treatment started on or after Day 8, most colonies could survive the treatment among dying fibroblasts, indicating a meaningful shift in reprogramming cell identity permissive to the small molecule treatment by that time (Figure 3C). Although early NIC&CTM treatment may prove useful, for the rest of this study the timing was performed as in Figure 1D and Figure 3F.

### Coordinated scRNA and *In-Vivo* Experiments

Given the transplanted iRPE data (Figure 2D,E,F), and the implications of NIC&CTM treatments (Figure 3, Figure S3), we designed an experiment to prepare iRPE from the same bulk passage with or without NIC&CTM treatment (iRPE/iRPENC). For each sample, the matured *BEST1::*EGFP positive cells were purified from one 6W by flow cytometry into 2 subsequent 24W wells for brief expansion and maturation with variable RPE-like stability (Figure 3D, Figure 3F). iRPENC treated cells maintained higher *BEST1::*EGFP expression thereafter, and with more lasting RPE-like ‘bleb’ that are common in high-quality iPSC.RPE cultures when cell junctions are tight and apical-basal flow bulges the RPE from the plate (Figure 3A, D, E; Video S3A, Video S3B). Unfortunately, several iRPENC ‘blebs’ had puckered and released from the plate to supernatant and were lost during media changes. For each, iRPE and iRPENC, one 24W well was sourced for subretinal transplantation into 4 immune-compromised albino rat retinas, and the other well was sourced for ~3500 cells in scRNA sequencing that also included standard Fib and iPSC.RPE culture cells prepared at the same time (Figure 3F).

### scRNA-sequence Profiling of iRPE System Including Starting and Target Cell States

The parallel cultures of Fib, iRPE, iRPENC, and iPSC.RPE were prepared for scRNA sequencing (Figure 3F). To prepare the reads for downstream analysis, we performed a workflow with several QC and read detection steps that allow for comprehensive uses in Seurat, Monocle, Velocyto, and SCENIC tools (Figure 4A) (Abugessaisa et al., 2020; Aibar et al., 2017; Butler et al., 2018; La Manno et al., 2018; Petukhov et al., 2018; Stuart et al., 2019; Trapnell et al., 2014).

**Figure 4:**
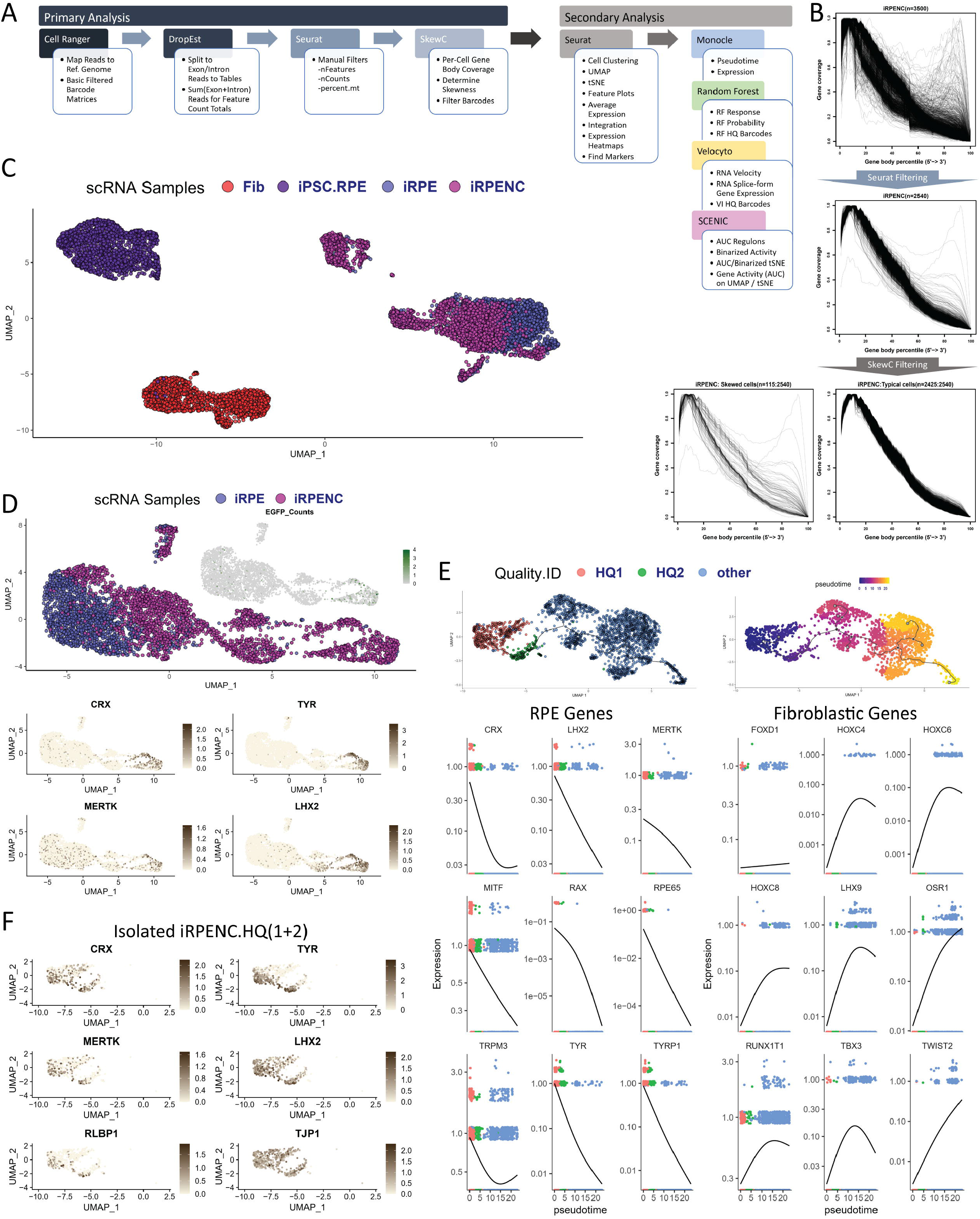
iRPE System scRNA Data Preparation and Analysis. **A)** An overview of the Primary and Secondary Analysis of scRNA samples in this study. **B)** Workflow 2 (serial) SkewC analysis total gene body mapped read traces, per cell after Cell Ranger (upper), after manual filtering in Seurat (mid) and then separated to Skewed cell traces (lower left) and Typical cell traces (lower right). **C)** Seurat UMAP plot of Fib, iRPE, iRPENC, and iPSC.RPE samples. **D)** Seurat UMAP plot of iRPE and iRPENC samples with inset coordinate plot of *BEST1::*EGFP counts (upper), and RPE gene expression feature plots (lower). **E)** Seurat UMAP coordinates of iRPENC plotted in monocle for HQ1, HQ2, and other, with pseudotime originating at the point of highest RPE gene and *BEST1:*EGFP expression (Figure S4D). Expression of RPE genes (left) and Fibroblastic genes (right) are plotted across pseudotime and labeled with HQ1, HQ2, and other. **F)** Seurat UMAP based RPE gene expression feature plots of iRPENC.HQ (HQ1 + HQ2).

### SkewC Improves Upon scRNA-sequenced Sample Distinction

To QC the resulting scRNA-seq data and decide the dataset for downstream analysis and interpretation, we employed workflows that combined commonplace manual filtering of poor quality cells and SkewC (Abugessaisa et al., 2020), a scRNA-seq quality assessment method. In short, SkewC uses gene body coverage of each single-cell to segregate skewed quality cells from typical cells based on the skewness of the gene body coverage.

From Cell Ranger filtered output, we performed two workflows, with manual filtering and SkewC filtering in parallel (Workflow 1; Figure S4A) or in series (Workflow 2; Figure 4B, Figure S4A). Samples were processed manually in Seurat (Butler et al., 2018; Stuart et al., 2019) with nFeatures, nCounts, and mitochondrial read frequency (percent.mt) for basic filtering parameters that output “Pass” or “Fail”; SkewC was implemented on the same cells with output “Typical” or “Skewed”. While manual and SkewC filtering both detected most low-quality cells, SkewC also frequently detected cells that clustered within the anticipated good cell clusters (Figure S4A, Figure S4B).

Interestingly, the Workflow 2 “Skewed” cells appeared to have normal counts and features and clustered accordingly. However, when we compared the feature average expression level for RPE marker genes, variable features, and all detected features, “Skewed” cells had a higher average expression than “Typical” cells, or no detection at all (Figure S4C). We believe SkewC possibly detected the single-cell libraries that had over-represented highly expressed reads and under-represented lowly expressed reads and were therefore possibly less dynamic. While both workflows improved filtering for clustering, Workflow 2 increased “Skewed” cell detection sensitivity within clusters (Figure S4A) and was used as the filter for the rest of the study (Figure 4B).

### iRPENC is Notably Improved and has ‘High-Quality’ Cells Approaching Subjective RPE Identity

Uniform Manifold Approximation and Projection (UMAP) (McInnes et al., 2018) analysis in Seurat clustered Fib, iPSC, and iRPE/iRPENC samples separately, but a distinct population closer to iPSC.RPE was predominately from iRPENC cells, suggesting that NIC&CTM treatments may have meaningfully improved iRPENC cells with lasting effect (Figure 4C). We therefore compared iRPE to iRPENC, and while many cells did cluster similarly in UMAP, the iRPENC mostly occupied the distinct population with *BEST1::*EGFP counts and enriched with important RPE features (*CRX, TYR, MERTK, LHX2*) (Figure 4D).

We then focused on iRPENC alone and performed Seurat clustering revealing clusters 1 and 4 to retain the highest expression of *BEST1::*EGFP and RPE features (Figure S4D); we labeled the iRPENC cells as HQ1 (cluster 1) and HQ2 (cluster 4), or ‘other’. We passed the UMAP coordinates to Monocle and employed Pseudotime analysis originating from the highest RPE feature rich region (Figure 4E). We found that, across Pseudotime, HQ1 and HQ2 cells tended to be high in RPE genes and low In Fibroblastic genes (Tomaru et al., 2014), while ‘other’ cells showed the opposite trend (Figure 4E). We thereafter combined HQ1 and HQ2, termed iRPENC.HQ, and the remaining cells as iRPENC.other. We were surprised that the distinction of iRPENC.HQ and iRPENC.other from the sample neatly showed that iRPENC.HQ had ‘percent.mt’ ratios matching iPSC.RPE cells while iRPENC.other had not (Figure S4E). Expectedly, iRPENC.HQ broadly retained important RPE gene expression (Figure 4F), while iRPENC.other did not (Figure S4F). Apparently, the iRPENC culture met a breaking point and diverged into a stable ‘HQ’ RPE-like population and a variably destabilized ‘other’ population with some donor cell (Fib) TFs.

### iRPENC and High-Quality Subset May Approach Objective RPE Identity

We sought to understand the identity of iRPENC cells objectively, and thus we employed Random Forest (RF) machine learning and RNA Velocity to do so (Breiman, 2001; La Manno et al., 2018).

With our Fib, iPSC.RPE, iRPENC.HQ, and iRPENC.other samples, we imported the public 5K Human PBMC dataset to increase the size and diversity of the RF and labeled 10 PBMC Seurat clusters (PBMC-CL0 to 9) (Figure 5A). The inclusion of more diverse cell types caused the iRPENC cells to cluster closer to the iPSC.RPE area of the UMAP (Figure 5A), while the Fib cells became more distant. We *trained* the RF on the PBMC, Fib, and iPSC.RPE samples, and then *tested* the iRPENC.HQ and iRPENC.other samples against that RF. RF Response is an absolute determination of singular identity and labeled most iRPENC cells with iPSC.RPE identity (Figure 5A, right). RF Probability of iRPENC was perhaps more informative, with high iPSC.RPE probability in iRPENC.HQ where Fib probability was often zero, and mediocre iPSC.RPE probability in iRPENC.other where Fib probability was more frequent (Figure 5A, lower).

**Figure 5:**
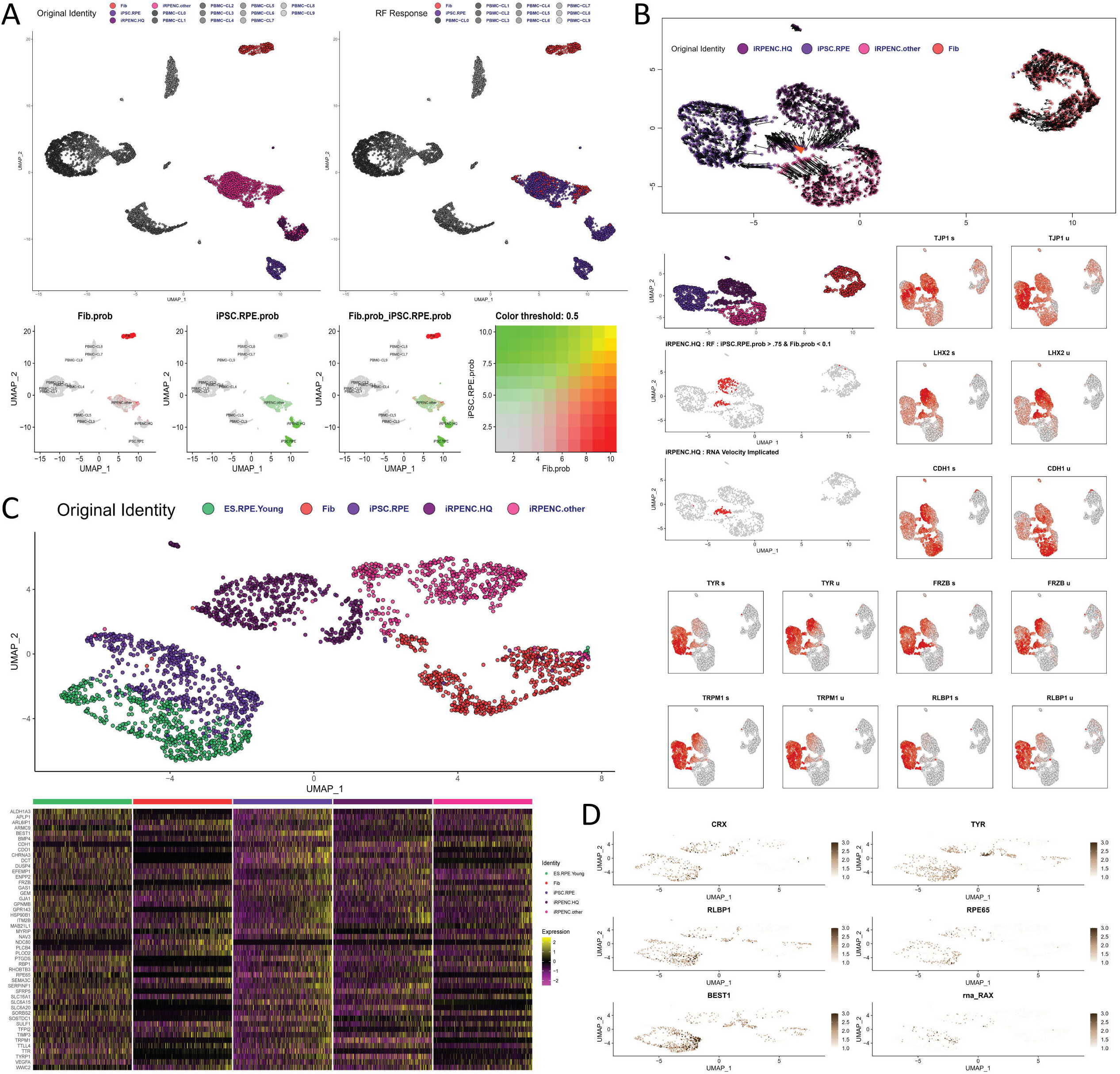
Machine Learning and Bioinformatics Interpretation of RPE Identity in iRPENC.HQ. **A)** Seurat UMAP plots of RF analysis samples labeled by Original Identity (upper left), RF Response (upper right), and by Fib/iPSC.RPE probability (lower). **B)** Seurat UMAP plots with cell velocity (upper), identity (mid), and RF / RNA Velocity Implicated RPE-like iRPENC.HQ cells indicated in the cell velocity map (orange arrow) and highlighted in red (mid). RPE gene expression detection by spliced/exon (s) and unspliced/intron (u) reads. **C)** Seurat UMAP plot of samples with reference based integration on iPSC.RPE cells (upper), including RPE gene heatmap (lower). **D)** The Seurat UMAP plot of Figure 5C with RPE gene feature plot for CRX, TYR, RLBP1, RPE65, BEST1, and RAX.

Since we could parse exon and intron counts from scRNA samples with DropEst (Petukhov et al., 2018), we prepared these matrices for RNA Velocity analysis with Velocyto (La Manno et al., 2018) on a Seurat UMAP with samples integrated on our reference iPSC.RPE (Figure 5B). Such integration placed the iRPENC cells much closer to the reference than without (Figure 5B). RNA Velocity showed that most clusters generally had arrows pointing inward, implicating a stable state. Surprisingly, a smaller group of iRPENC.HQ had significant RNA Velocity in the direction of iPSC.RPE, implicating state change toward iPSC.RPE state (Figure 5B). We therefore specifically labeled those ‘RNA Velocity Implicated’ cells on the UMAP, and found that they neatly overlapped with 51 of the highest RF probability RPE-like cells of iRPENC (iPSC.RPE.prob >75%, Fib.prob <10%) (Figure 5B). Taken together with recurring cell clustering trends, we concluded that unique machine learning and bioinformatics tools could find agreement among highly dynamic and specific criteria among unique cell systems.

We further observed from the unspliced (u) and spliced (s) Veloctyo plots of important RPE genes, that some RPE RNAs (*CDH1, FRZB, LHX2, TJP1*) were expressed typically in iRPENC.HQ as in iPSC.RPE, while other RPE genes (*RLBP1, TRPM1, TYR*) showed splice variance that related important epigenetic regulation was involved, and that the ‘RNA Velocity Implicated’ cells were perhaps the most like iPSC.RPE (Figure 4B).

Integrating on iPSC.RPE as reference brought iRPENC and its iRPENC.HQ closer to iPSC.RPE in UMAP (Figure 5B), with RPE/RPE-like cells overlapping in principal component space (Figure S5A). However, we sought to include external RPE data to improve our analyses. We imported a count matrix from a recently published ES cell-derived RPE study (Lidgerwood et al., 2019) as ‘ES.RPE.Young’ and integrated the data with iPSC.RPE reference. We found that ES.RPE.Young intermixed with iPSC.RPE and our iRPE system trend of reprogramming from Fib toward RPE became more apparent (Figure 5C). While positions across UMAP_2 appeared dynamic, positions across UMAP_1 appeared to better reflect the spectrum across Fib and RPE identities. The trend was not obvious by simpler principal component space where the ES.RPE.Young, iPSC.RPE, and iRPENC.HQ shared the same general area while Fib and iRPENC.other were usually further away (Figure S5B). A heatmap of gene expression based on specific molecular signatures of primary RPE cells and clinical iPSC.RPE cells (Kamao et al., 2014; Liao et al., 2010) showed that ES.RPE.Young, iPSC.RPE, and iRPENC.HQ had a shared pattern, while Fib and iRPENC.other also had a different shared pattern (Figure 5C, lower), supporting the trends seen in the Pseudotime gene expression plots (Figure 4E). To confirm that observation, we then checked Seurat expression for (*CRX, TYR, RLBP1, RPE65, BEST1, RAX*), and found relatively comparable expression between the ES.RPE.Young, iPSC.RPE, and iRPENC.HQ cells (Figure 5D).

### Gene Regulatory Network Analysis Reveals Distinct Cell Type Signatures

To better understand the reprogramming status of iRPENC cells, we employed SCENIC (Aibar et al., 2017) to determine upstream ‘regulons’ from putative downstream expression at the single-cell level yet often analyzed in specified clusters (Figure S6A). We compared Fib, iPSC.RPE, iRPENC.HQ, and iRPENC.other at equal cell numbers per sample and performed tSNE in Seurat (Figure S6B). Global AUC per sample indicated iPSC.RPE AUC patterns resembled iRPENC.HQ (Figure S6C) despite a notable difference at the few lowest row-clusters where differences were obvious.

Many TFs operate in OFF/ON states relative to dosage and cofactor availability. In SCENIC analysis, manually thresholding AUC histograms for binarization (OFF/ON) of the regulons provides an easier to interpret binarized plot (Figure 6A, left) than without (Figure S6D). Still, in all cases, the iPSC.RPE clustered with iRPENC, and more closely to iRPENC.HQ (Figure 6A, S6D). A distinct large set of Fib regulons was effectively OFF in the iPSC.RPE and iRPENC.HQ samples (Figure 6A, Figure S6A). Among the top 281 regulons, the pattern of iPSC.RPE and iRPENC.HQ regulon activity was similar in the higher and lower activity regulons (Figure 6A, left). To succinctly understand the most relevant data, and the cell reprogramming, we reduced the plot to the Top 55 Regulons affecting >95% of cells in the analysis (Figure 6A, mid) and saw that most Fib regulons were OFF in iRPENC.HQ where most iPSC.RPE regulons were ON. Not surprisingly, the iRPENC samples also had a specific subset of regulons whose expression likely restricts iRPENCs from closer clustering to RPEs. Of interest, the MITF and CRX regulons were strong in iRPENC.HQ, as in iPSC.RPE, indicating that the removal of conditional reprogramming left these endogenous regulatory networks intact (Figure 6A). However, the SOX9 and PAX6_extended regulons were very poorly represented in iRPENC.HQ restating the potential role for these factors in our iRPE system despite the antagonistic effects when expressed from Day 0. Expectedly, checking average gene expression for the TFs of the same regulons correlated the relationship between most TF gene expression and putative downstream regulatory activity (Figure 6A, right).

**Figure 6:**
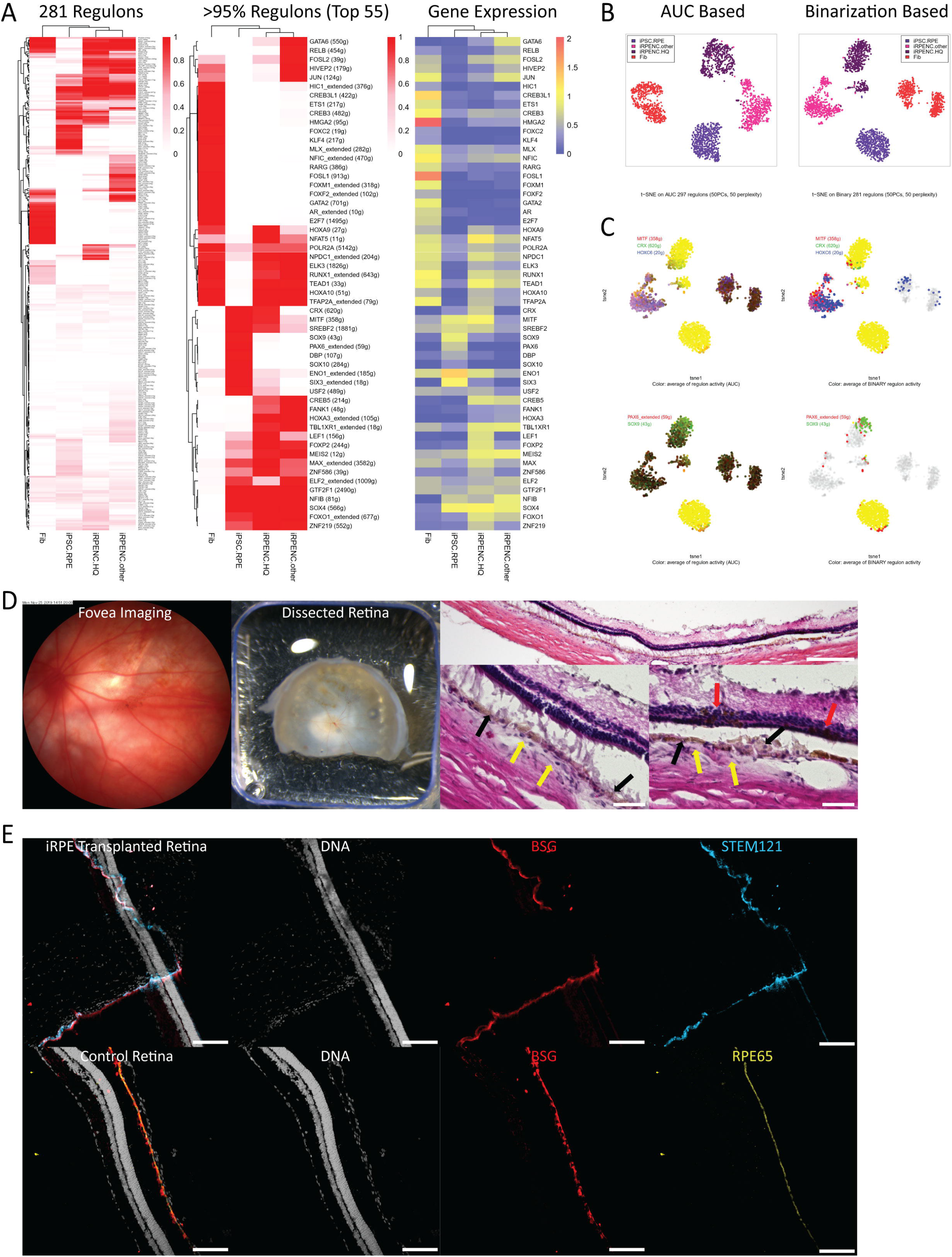
iRPE System Genetic Regulatory Analysis and Validation *In-Vivo*. **A)** SCENIC Binarized regulons total (281, left), top 55 (mid) with heatmap(pheatmap) clustering and regulon activity (red scale), with row-matched TF expression (log-transformed TF gene expression averages per sample) (right). **B)** SCENIC AUC Based (left) and Binarization Based (right) tSNE clusters of samples. **C)** AUC (left) and Binarized (right) activity of MITF, CRX, HOXC6, PAX6_extended, and SOX9 regulons. **D)** Live fovea imaging of a iRPENC transplanted albino rat retina (left), with apparent pigmentation during dissection (mid). Cryosection H&E stains (right, low mag (upper) high mag (lower x2)) show pigmented RPE-like cells (black arrows) in expected areas sometimes interfacing photoreceptor outer segments. *Note: A non-pigmented RPE layer is indicated (yellow arrows), and out-of-place pigmentation is also noted (red arrows). Upper Scale bar = 200μm, Lower Scale bars = 50μm.* **E)** A proximal cryosection (upper) to Figure 6D H&E stains was used for IHC against BSG (red) and STEM121 (light blue), with DNA counterstain (light grey). An untreated control retina (lower) was used for comparable IHC against BSG (red) and RPE65 (yellow), with DNA counterstain (light grey). *Scale bars = 100μm.*

On these samples, we performed a Seurat UMAP plot (Figure S6E), and tSNE plots based on AUC (Figure 6B, left) and Binarized Regulons (Figure 6B, right). Expectedly, UMAP provided the distinct clustering that informed sample differences (Figure 6E), and as reported in SCENIC (Aibar et al., 2017) the AUC based and Binarization based tSNE plots improved cell clustering more than when tSNE is based on gene expression (Figure 6B, Figure S6B,E). Curiously, in these plots, and others in this report, a few iPSC.RPE plot neatly among the iRPENC.HQ cluster.

To further understand our cells, we selected the Binarization based tSNE plot to coordinate AUC and Binary Regulon activity of MITF, CRX, and HOXC6 regulons. Expectedly, iRPENC.HQ and iPSC.RPE showed similar RPE regulons (MITF, CRX) while iRPENC.other and Fib showed the HOXC6 fibroblastic regulon (Figure 6C). Binarization thresholding can be variable, for example we set the CRX threshold to the highest of three normal distributions of AUC activity where the iRPENC.HQ and iPSC.RPE were comparable and mixed, yet the middle distribution represented iRPENC.other cells, and the lowest was Fib cells (Figure S6F). We also looked at candidate iRPE factors PAX6_extended and SOX9 (Figure 6C, lower, Figure 2A), which interestingly showed that a few iRPENC cells strongly represented the PAX6_extended regulon, and a cluster of iRPENC.HQ had near full SOX9 regulon activity. We then plotted the individual AUC plots for HOXC6, CRX, MITF, PAX6_extended, and SOX9 regulons across the UMAP and tSNE coordinates to clarify local enrichment (Figure S6G). Interestingly, plotting the ‘RNA Velocity Implicated’ iRPENC.HQ cells on the Binarization based tSNE coordinates showed that the separated subset of cells toward iPSC.RPE were predominately those with RNA Velocity in that same identity direction (Figure S6H, Figure 5B). Taken together, gene regulatory network inference for cell identities from SCENIC and cell state change inference from RNA Velocity/splicing (Velocyto), could unify to implicate an ideal subset of cells for focused analysis.

### scRNA Sampled iRPENC Survived and Integrated to Host Retina *In-Vivo*

Parallel cultures of iRPENC used for scRNA were also transplanted to immune compromised albino rat retinas (Figure 3F, Figure 6D). Transplanted iRPENC had apparent pigmentation noted in standard fovea imaging just 6 weeks after transplant, and with obvious pigmented tissue in the retina about 5 months after dissection for cryosection and immunohistochemistry (IHC) (Figure 6D). H&E stains of transplanted retinas showed pigmented cobblestone cells in many areas, particularly interfacing with some host photoreceptor outer segments (Figure 6D). Transplanted iRPENC cells appeared to sit atop host albino RPEs, with notably similar morphology and H&E stain characteristics. In some cases, RPE-like cells with weak pigment speckles were noted atop the retina ganglion cell layer, which can happen when transplant areas leak cells to the vitreous. Curiously, some pigment, or pigmented cells, appeared at the outer nuclear layer *membrana limitans externa*, which piqued concern for material transfer or mis-localization between the pigmented iRPENC and host cells (Figure 6D).

To validate the RPE positioning and polarity, we then prepared IHC of intervening cryosections proximal to the H&E stained sections in Figure 6D. We found that the apical RPE marker protein BSG (Deora et al., 2004) had labeled the apical face of the RPE cell layer (Figure 6E). Expectedly, many BSG+ cells co-localized apical to STEM121, a common human-specific cytosolic marker (Tu et al., 2019) (Figure 6E). To validate our apical BSG protein detection in host RPE cells and IHC methods, we stained untreated control retina cryosections with BSG and the RPE marker RPE65 (Figure 6D).

Taken together, these observations strengthen the notion that the optimized iRPE reprogramming system conditions may reprogram human somatic cells into stabilized cells with subjective and objective metrics for RPE identity that could mature, integrate, and interface in transplanted host retinas.

## DISCUSSION

### Bulking Up

In basic research we can explore nature willy-nilly, yet medicine is bound by economies of scale and practical finance. The adage ‘*time is money’* holds true in medicine, inspiring research for shortcuts toward autologous or compatible cell therapies. In that context, individual colonies of iRPE were irrelevant and only bulk cultures proved efficient with robust expansion toward the excess, then purity, that is often necessary in biomedical products. In bulk, many stabilized iRPE cells could be found after ~2 months’ time which suggests significant savings in contrast with iPSC.RPE generation that requires > 6 months. To our surprise, Nicotinamide and Chetomin provided an unforeseen role in iRPE cell selection, although a complete purification of high-quality iRPEs remains unaddressed. For that reason, we foresee necessary co-development of the iRPE reprogramming system with plausible cell purification technologies (Miki et al., 2015; Ota et al., 2018; Plaza Reyes et al., 2020). Relatedly, earlier iRPE may prove better given the fact that FACS purified *BEST1::*EGFP+ iRPENC cells later divided into ‘HQ’ and ‘other’ in time, and that several studies find that overly-mature RPEs are poor candidates for cell therapy.

### Cell Reprogramming

This iRPE system may advance timely somatic cell conversion toward RPE given the hallmark of stability after the removal of conditional reprogramming; a state not shown in other iRPE systems (D’Alessio et al., 2015; Rackham et al., 2016; Zhang et al., 2014). We recognize the importance of previous reports variably employing MITF, OTX2, and CRX in wet cultures to induce RPE features (D’Alessio et al., 2015; Zhang et al., 2014). Still, prior iRPE reports ostensibly showed ‘direct reprogramming’ while relying on the pluripotency factor OTX2 (Buecker et al., 2014; Thomson and Yu, 2012) and optionally relying on the Yamanaka factor KLF4 (Zhang et al., 2014). Given the state of the cell reprogramming field, ‘direct reprogramming’ may be a misnomer since most reprogramming system cell state changes, over significant time, is diverse. We wonder how a slow/non-dividing somatic state could generate another slow/non-diving somatic state, effectively, without the proliferative precursor programs that predicate important *survival* and *generative* identity. Conversely, this study sought to exploit that hypothesis, focusing on plasticity, precursor, lineage, and end-state. Given that iPSC-like reprogramming MET was evident in the first week of iRPE reprogramming, and cell death/survival was obvious, we anticipate that tumor suppressors like p53 or Rb may perturb iRPE reprogramming, or selectively affect surviving cell states and genomic stability (Kareta et al., 2015; Marión et al., 2009; Samavarchi-Tehrani et al., 2010).

Among the TFs in this iRPE system, Microphthalmia-Associated Transcription Factor (MITF) is a regulator of RPE differentiation (Adijanto et al., 2012; Hansson et al., 2015) that critically cooperates with Orthodenticle homeobox 2 (OTX2) for RPE development (Bharti et al., 2006, 2012; Ramón Martínez-Morales et al., 2004). Both MITF and OTX2 are pioneering TFs (Soufi et al., 2015), and OTX2 is developmentally retained from primed pluripotency as an organizer/specifier/reprogrammer (Buecker et al., 2014; Shahbazi et al., 2017; Thomson and Yu, 2012), through the neural plate, optic vesicle, and RPE specification (Hever et al., 2006). LIN28 and MYCL, the ‘hUL’ cassette for iPSC reprogramming enhancement (Okita et al., 2011), are also involved in natural retinal regenerating reprogramming (Luz-Madrigal et al., 2014). LIN28 binds and neutralizes *Let-7*, a promiscuous and broadly expressed miRNA somatic cell identity ‘lock’ against plasticity, reprogramming, and state change. MYCL is a form of MYC, which cooperates with reprogramming TFs by holding open freshly pioneered nucleosome-bound genomic DNA (Soufi et al., 2015). The combination (MITF, OTX2, LIN28, MYCL) was improved by CRX, a powerful Orthodenticle homeobox TF involved in various aspects of retinal differentiation and early RPE fate (Esumi et al., 2009; Furukawa et al., 1997).

CRX was important to this iRPE system and was identified by SCENIC as a distinguishing regulon for iPSC.RPE and iRPENC.HQ cells. However, CRX is lowly expressed or turned off *in-vivo* as RPE matures, highlighting a potential role in regenerative medicine. Comparable MITF and CRX regulons in iPSC.RPE and iRPENC.HQ, among absence OTX2 regulon relevance, provides and strong backdrop for interpreting necessary transient, or lasting, reprogramming TFs; indeed, most iRPENCs did not express detectable endogenous OTX2. Weak iRPENC.HQ signatures for PAX6_extended and SOX9 regulons reiterate their value in other iRPE systems; these TFs could not be used from Day 0 in our iRPE system context, and thus an exploration of differential timing, or other TF candidates, may significantly improve iRPENCs to match the iPSC.RPE model. Importantly, the current iRPENC.HQs stability may exist due to the loss Fib-specific regulon activity. With stability, perhaps most of the reprogramming was achieved, leaving a fraction of meaningful donor cell gene regulatory networks to address, and only if they may affect the safety or function of the target autologous RPE cell product.

### Cell Identity

iRPENC displayed numerous characteristics of RPE *in-vitro*, and *in-vivo* when transplanted to immune compromised albino rat retinas where cells survived, expressed a maturation reporter, pigmented, and apparently integrated into host RPE layers sometimes interfacing with host photoreceptor outer segments. The scRNA analysis of iRPEs revealed a distinct sub-population of ‘high-quality’ cells that were better stabilized toward RPE cell identity and whose generation was drastically improved by the addition of Nicotinamide and Chetomin during cell reprogramming. Both machine learning Random Forest and RNA Velocity approaches objectively strengthened that subjective segregation and further identified subsets of iRPEs with high RPE expression and genetic regulation. Taken together, the bioinformatics tools employed here provided a window for analysis that helped us characterize and understand the iRPE system and important cell identities, while providing extensive data that we have yet to fully explore.

Of note, the inclusion of other cell samples, and cell numbers, strongly affected bioinformatics analysis. In Random Forest, more distant blood cell types caused iRPENC to cluster closer to iPSC.RPE in UMAP and strengthened the RF Response and Probability clarity between Fib and iPSC.RPE identities. In Veloctyo, the absence of other cells brought iPSC.RPE and iRPENCs yet closer. Conversely, when the ES cell-derived RPE model was included, distance increased between the iRPENC.HQ and iPSC.RPE UMAP space despite overlapping principal component space coverage. Even adjustments to cell numbers often had notable effects on the positioning and clustering of cells. For these reasons, careful per-sample consideration may be critical for scRNA analysis.

## EXPERIMENTAL PROCEDURES

### Animal Use

Rat handling and transplant experiments were carried out with humane methods in compliance with animal ethical standards approved by RIKEN Kobe Safety Center.

### Human Fibroblast Culture

BJ Human Foreskin Fibroblasts (ATCC), were cultured in fibroblast culture media (FCM) that consisted of 90% DMEM-high glucose, 10% Tetracycline free Fetal Bovine Serum (FBS), and 1% Penicillin Streptomycin (P/S) with no filtering. For most cultures, 10cm cell culture plates were coated with 0.1% gelatin in CMF-DPBS for 1 hour. Cryopreserved fibroblasts were thawed, diluted with FCM, and centrifuged at 200 x g for 4 min, and then the pellet was resuspended for cell counting and then diluted appropriately in FCM to yield ~650,000 cells/10mL, triturated to mix evenly. The gelatin/PBS coating was aspirated from the 10cm plate and 10mL of diluted cells in FCM were plated. Cells were incubated at 37°C until ~ 85-90% confluent and then passaged or used for reprogramming experiments.

### iRPE Reprogramming

iRPE Media consisted of 85% DMEM/F12+GlutaMax (1X), 15% KnockOut Serum Replacement, 1% MEM Non-Essential Amino Acids, 1% N-2 Supplement (all filtered via a 0.22μm polyethersulfone – PES) and then prepared as frozen aliquots. 1% Penicillin Streptomycin (P/S) is added fresh, and in the first 10 days of reprogramming Basic Fibroblast Growth Factor (bFGF) is added to a final concentration of at 10ng/mL with 2-mercaptoethanol (2ME) at 1:1000 dilution. Doxycycline is added at 1μg/mL (See Figure 1) for the Phase 1 media. Phase 2 media consisted of all the above-mentioned reagents except for bFGF and 2ME.

Generally, a 10cm plate was coated with 0.1% gelatin and plated with 650,000 fibroblasts containing conditional doxycycling-inducble reprogramming sets we generally designated as ‘programs’ (e.g. Program 2, Set 2 = P2.2). Fibroblasts ready to reprogram were incubated 37°C until ~ 85-90% confluent. Fibroblasts were reprogrammed in 6W plates. Briefly, target wells of 6W plates were coated with 1.5 mL of 1:150 iMatrix511/CMF-DPBS substrate for 1 hour at room temperature. Fibroblasts were passaged to yield ~125,000 cells/1.5 mL per well, and then incubated at 37°C 16-24 hours before reprogramming medium (Phase 1 iRPE medium) was added. Fibroblasts were checked prior to reprogramming to ensure that the cells plated as single evenly dispersed cells. The addition of the reprogramming medium marks the beginning of the reprogramming or conversion process and is noted as Day 0. The Phase 1 medium is added fresh at 2mL every day together with any molecule or supplement for 9 more days – Day 0 to Day 9, which makes 10 continuous days of feeding the cells with the Phase 1 media. On the Day 10, the Phase 1 medium was replaced with Phase 2 medium (no bFGF and no 2ME) at 2mL every-other-day.

Phase 2 media was used until ~Day 28-32 when the cells were passed in bulk, combining all colonies, to an iMatrix511 coated 6W plate. Cell passage was as described with fibroblasts, but with Phase 2 medium containing 10% FBS to neutralize trypsin. In some cases, defined trypsin inhibitor was used. Reprogramming cells were plating between 300,000 to 500,000 cells/well of a 6W plate in Phase 2 medium and incubated at 37°C until for ~48 hours before refreshing the Phase 2 medium. On ~Day 36 Phase 2 medium was prepared with fresh Nicotinamide [5-10mM] and Chetomin [40-80nM] (NC) and fed every other day for 10 days. Cells that did not receive NC, were fed the same medias excluding those molecules. After the last feed with NC, the reprogramming cells were fed Phase 2 medium once more, before culture in Maturation Medium ~Day 50. *Note: If necessary, colonies were counted on Day 9 and the diameter of the colonies were measured on Day 13.*

### iRPE/iPSC.RPE Maturation Media Culture (SFRM-B27)

Maturation Media contained 70% DMEM-low glucose, 30% Nutrient Mixture F-12 Ham, 1% GlutaMax (100X), 2% B-27 Supplement (all filtered via a 0.22μm polyethersulfone – PES) and stored in frozen aliquots. Fresh 1% P/S, 0.5-1μM SB431542, and 10ng/mL bFGF is added before use.

On ~Day 50 of iRPE reprogramming, the medium is changed to the Maturation Media (SFRM-B27) with 1μg/mL Doxycycline. The Doxycycline concentration is gradually reduced from 1μg/mL for the first and second feeds, and then 0.5μg/mL for the next two feeds. From the fifth feed with Maturation Media, doxycycline was not added. iRPE or iPSC.RPE were fed Maturation Media every-other-day, as necessary. *Note: iPSC.RPE could be fed Maturation Media with or without doxycycline without consequence since they did not have doxycycline-inducible reprogramming factors.*

### Single-cell RNA Sampling

Cells were dissociated and passed through cell screen cuvettes to isolate mostly healthy single-cells that were prepared with 10x Chromium Single Cell 5’ Library & Gel Bead Kit (PN-1000014) skipping steps 4/5 per standard protocol for non-VDJ samples. Sample libraries were finalized and sequenced on one Hiseq X lane (150bp PE; Macrogen) for each. Standard Cell Ranger protocol detected sample chemistry and produced ‘possorted’ BAM files from which the subsequent Primary Analysis workflow in Figure 4A was performed.

### Albino Rat Xenografts of iRPE/iRPENC

Briefly, cells for transplant were prepared similar to cell passage, but kept on ice, and at 50,000 cells per μL. Recipient rats were prepared with isofluorane and accommodated, and then 1-2μL of cells added to the subretinal space between the neural retina and RPE/Choroid. Rat health was checked frequently, and transplanted animals were generally maintained for 4-5 months prior to sacrifice.

### Sample Fixation and Cryosection

Rats were sacrifice humanely, and the eyes extracted and fixed with 4% paraformaldehyde in PBS solution, overnight. The eyes were then desiccated in 30% sucrose solution for ~24-48 hours. Upon later dissection, based on prior fundus imaging, the exact pigmented or transplanted parts of the eye were retained, and the rest discarded. Samples were placed in a 10mm X 10mm X 5mm size Tissue cryomold and filled with optimal cutting temperature (O.C.T.) compound, and then stored at −80°C.

For cryosection, the frozen cryomold containing the eye tissue was sectioned with a Thermo Scientific Micron HH 560 cryomictrotome. Each section had a fine size of 12micron, placed on glass slides, and then stored at −80°C until further use.

### H&E Staining

Generally, for each eye sample, every fifth cryosection slide was air dried for 1 hour, and then subjected to common H&E stains procedures.

Briefly, residual O.C.T. compound was dissolved by immersing in CMF-PBS for 1 hour. The slides were then placed in Hematoxylin stain for 2.5mins, followed by two MilliQ water washes of 2.5 minutes. The slides were then placed in Eosin stain for 1 minute, followed by MilliQ water washes for 1-2 seconds. The slides are then moved to 80% ethanol, then 90% ethanol washes for 30 seconds each. Then, slides pass through two 100% ethanol washes for 3 minutes each. Finally, the slides pass through two washes in xylene for 3 minutes each. After the xylene, the slides are air dried and then mounted with Malinol and nail polish prior to imaging.

### Immunohistochemistry and Confocal Microscopy

Immunohistochemistry in Figure 6E was performed on previously noted cryosections. Sample slides were removed from the freezer and then air dried at room temperature for ~1 hour. Slides were submerged in 100°C 1X citrate buffer for 20 mins for heat induced epitope retrieval. Generally, blocking buffer (BB) was 4% horse serum in CMF-DPBS. Samples were permeabilized for 30 minutes with 0.2% Triton X-100 in BB, and then blocked for ~1 hour in BB. Primary antibody solutions were prepared in BB and then added to slides and incubated overnight at 4☐°C.

All samples were washed with BB 3 times, followed by secondary antibody solution for ~1.5 hours at room temperature. The solution is replaced with DNA stain (Hoechst 33342 1:2000) for ~10 minutes. The slides are washed two more times with CMF-DPBS, and then dried and mounted FluorSave and a coverslip prior to imaging with Zeiss LSM 700 Confocal Microscope.

#### Antibodies Used

| Reagent | Maker / Catalog # | Dilution |
| --- | --- | --- |
| Mouse anti-RPE65 | Merk Millipore MAB5428 | 1:250 |
| Goat anti-BSG (CD147/EMMPRIN) | R&D Systems AF972 | 1:20-40 |
| Mouse anti-STEM121 | Takara Bio Y40410 | 1:250 |
| Donkey anti-Mouse Alexa Fluor 647 | Thermo Fisher A-31571 | 1:1000 |
| Donkey anti-Goat Alexa Fluor 546 | Thermo Fisher A-11056 | 1:1000 |
| Hoechst 33342 | Thermo Fisher H3570 | 1:2000 |

## Supporting information

Supplemental Figures and Videos (see MS for legends)

## AUTHOR CONTRIBUTIONS

Conceptualization, C.K.; Methodology, C.K. and I.N.W; Cryosections, J.S., M.N., H.H. and H.S.; Formal Analysis, C.K. and I.N.W.; Investigation, C.K., I.N.W., I.A., B.K, HH.C., and M.M.; Resources, C.K., A.T., J.S., I.A., B.K., P.C., E.A., Mi.T., Y.S., M.M. and Ma.T.; Writing – Original Draft, C.K.; Writing – Revision & Editing, C.K., I.N.W., A.M., P.C., I.A., and E.A.; Visualization, C.K., H.S., M.N., and I.N.W.; Project Supervision, C.K., and Ma.T.; Retina Aspect Supervision: M.M., A.M., and Ma.T.; Bioinformatics Aspect Supervision: E.A., I.A., T.K., B.K., and P.C.; Project Administration, C.K. and Ma.T.; Funding Acquisition, C.K., M.M., and Ma.T.

## ACKNOWLEDGEMENTS

We honor the help and support members of the RIKEN Lab for Retinal Regeneration with special mention for Sunao Sugita. We are grateful for Grace Lidgerwood & Alice Pébay of the University of Melbourne, Australia, and Anne Senabouth & Joseph Powell of the Garvan Weizmann Centre for Cellular Genomics, Australia, for sharing their ES cell-derived RPE scRNA data tables for use in our study. We greatly appreciate the help of Teruaki Kitakura of RIKEN Center for Integrative Medical Sciences (DGM), Japan, for technical support and setting the computing environment. We also thank Osamu Nishimura and the Shigehiro Kuraku Lab at RIKEN for providing a HPC for initial Cell Ranger processing.

## SUPPLEMENTAL FIGURE LEGENDS

**Figure S1: Related to Figure 1 & Figure 2**

**A)** A representative dox-inducible system viral construct with tetracycline (doxycycline) response elements upstream of an example TF/exogene, MITF.

**B)** A picked and sub-cultured pigmented iRPE colony after 100 days of reprogramming (left), was passaged for expansion (mid) and developed into large pigmented cells resembling aged RPE based on size, polynuclei, and morphology (right). *Scale bars = 500μm.*

**C)** A table of the initial reprogramming factor virus set in the parental culture (e.g., MITF, OTX2, LIN28, MYCL, CRX) and the Genomic DNA PCR detection (YES/NO) for those virii in individually sub-cultured/cloned iRPE colonies.

**Figure S2: Related to Figure 2**

**A)** An iRPE transplanted albino rat retina cryosection with H&E staining to reveal pigmented human iRPE cells (black arrows) in ‘rosettes’ in the neural retinal space.

**Figure S3: Related to Figure 3**

**A)** PEDF (left) and VEGF (mid) secretion ELISA detection from transwell Apical (upper) and Basal (lower) cell supernatants of hRPE, iRPE, and iRPE+CTM treated cells, sampled after Week 1 and Week 4. TER for the same cultures was taken weekly and plotted (right).

**B)** A bulk iRPE passage was prepared and treated with NIC, CTM, and NIC&CTM, or control. *BEST1::*EGFP expression during treatment is represented (left) and pigmented morphological imaging after treatment, in maturation, is represented (right). *Left scale bars = 100μm, Right scale bars = 500μm.*

**Figure S4: Related to Figure 4**

**A)** Seurat UMAP plots of Manual Filtering and SkewC labeled or filtered cells in Workflow 1 (parallel; upper), and Workflow 2 (series; lower).

**B)** SkewC cell clustering by gene body coverage with ‘Typical cells highlighted in red.

**C)** Box plots of the log-transformed Feature Average expression levels for all samples in Figure S4A, based on feature sets of RPE Marker Genes, Seurat Variable Features, and all Features. Skewed cell ‘no detection’ is indicated with red arrow.

**D)** Seurat UMAP plot of iRPENC cell clusters (upper left), *BEST1::*EGFP expression counts (upper right), and monocle gene expression for CRX, TYR, MERTK, and LHX2 plotted on the Seurat UMAP coordinates.

**E)** Seurat violin plots for calculated ‘percent.mt’ for Fib, iPSC.RPE, iRPENC.HQ, and iRPENC.other samples.

**F)** Seurat UMAP based RPE gene expression feature plots of iRPENC.other.

**Figure S5: Related to Figure 5**

**A)** Seurat Principal Component (PC1 vs PC2) plots of cells analyzed in Figure 5B.

**B)** Seurat Principal Component (PC1 vs PC2) plots of cells analyzed in Figure 5C.

**Figure S6: Related to Figure 5 & Figure 6**

**A)** SCENIC boxplots for the number of cells per regulon, and the number of regulons per cell.

**B)** Seurat tSNE plot for the samples used in SCENIC analysis.

**C)** SCENIC total AUC regulon activity.

**D)** Heatmap (pheatmap) with clustering of all AUC regulon activity per sample type.

**E)** Seurat UMAP plot for the samples used in SCENIC analysis.

**F)** Example ‘Binarization’ AUC threshold setting for CRX regulon, indicating the OFF and ON determination.

**G)** Gene set AUC for HOXC6, CRX, MITF, PAX6_extended, and SOX9, plotted across Seurat UMAP coordinates (upper) and SCENIC Binarized tSNE (lower) coordinates.

**H)** Seurat tSNE plot using SCENIC Binarized tSNE coordinates to highlight RNA Velocity Implicated iRPENC.HQ cells.

**Video S3: Related to Figure 3**

**A)** A representative video of a slow change of focal (Z) position of iRPE pigmented ‘bleb’ cells and attached culture.

**B)** A representative video, matching Video S3A, imaged for *BEST1::*EGFP.

/end/

