## Supplementary figures and images for "Inducing Human Retinal Pigment Epithelium-like Cells from Somatic Tissue"

### Figure S1.png

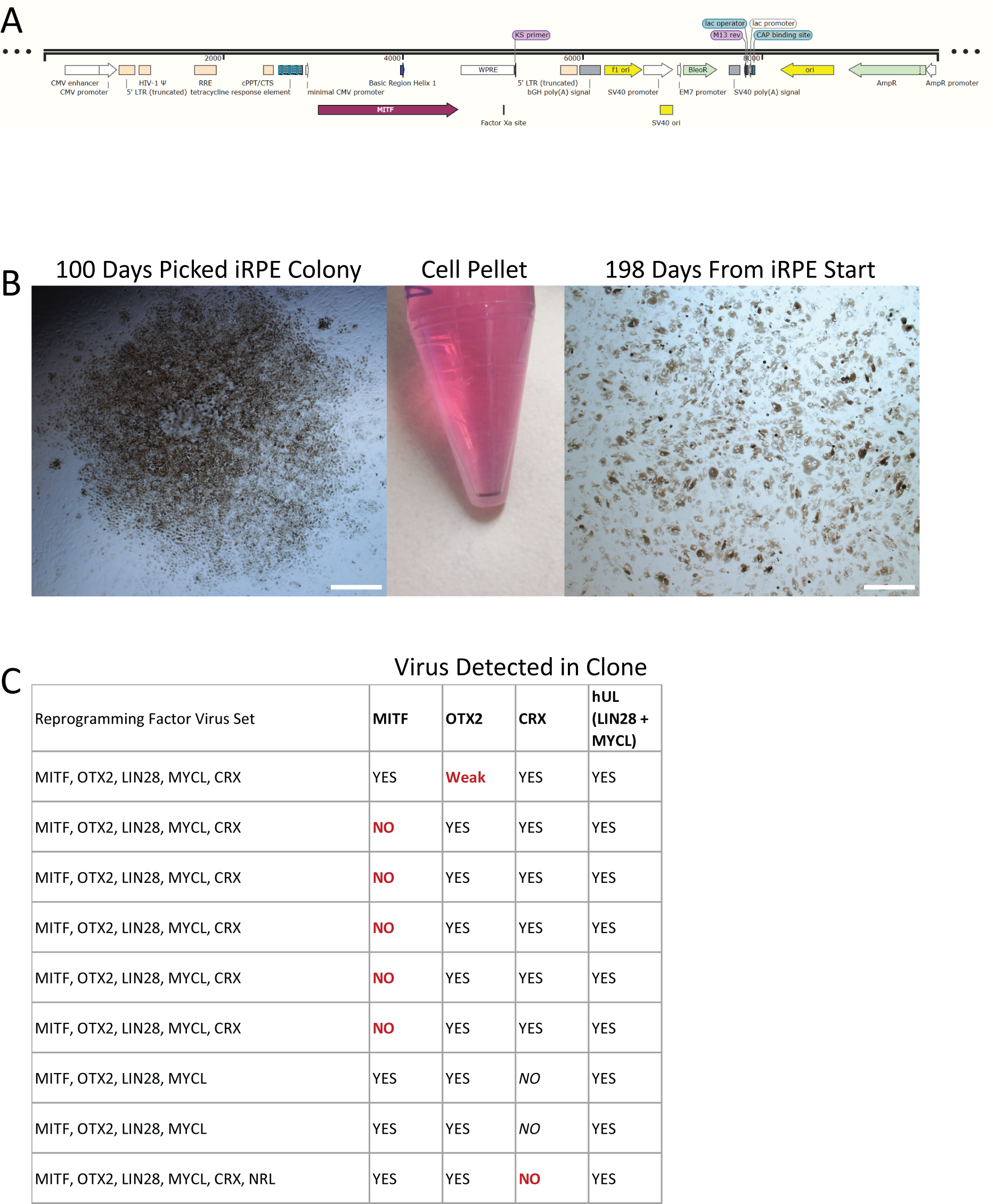

### Figure S2.png

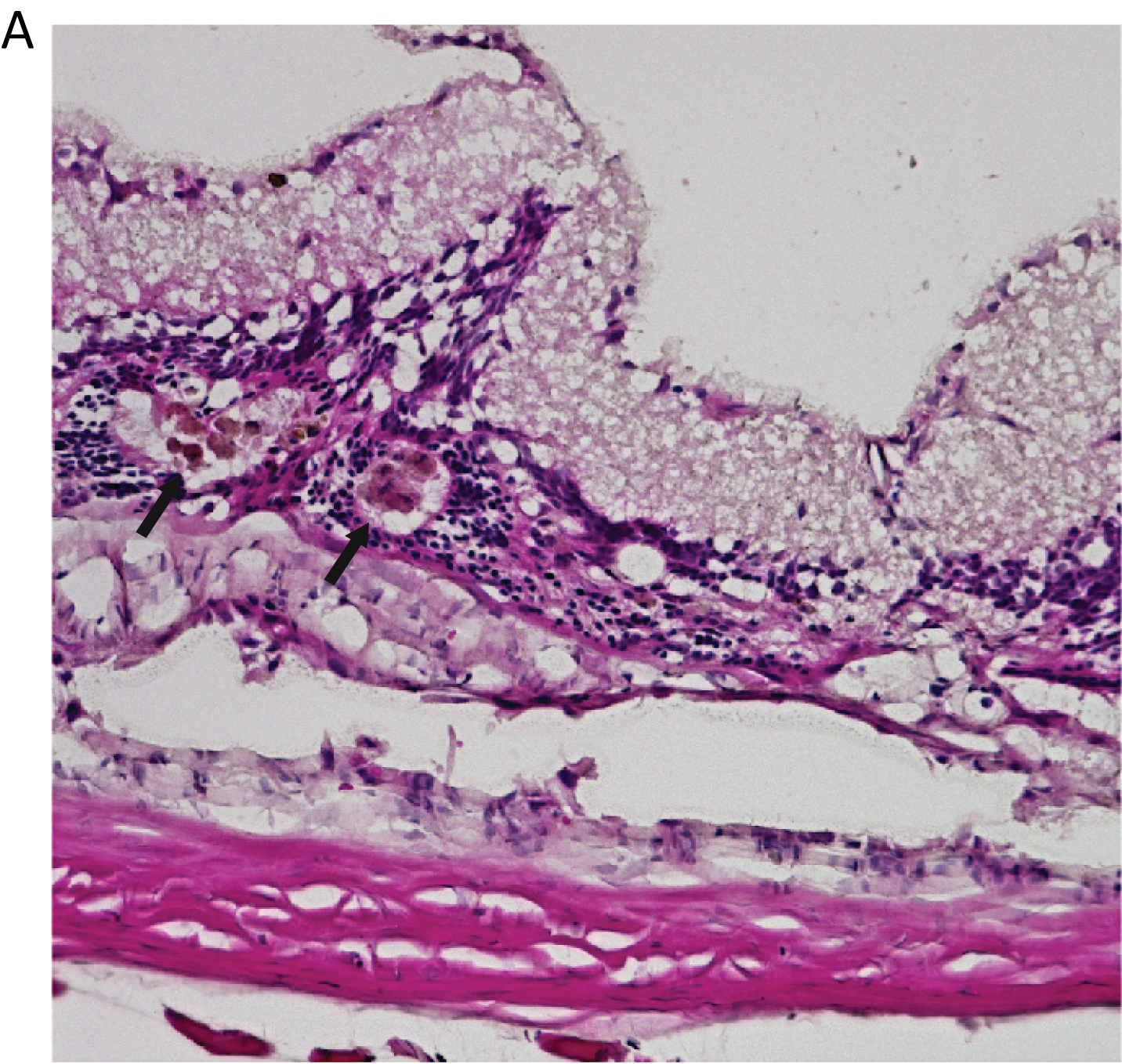

### Figure S3.png

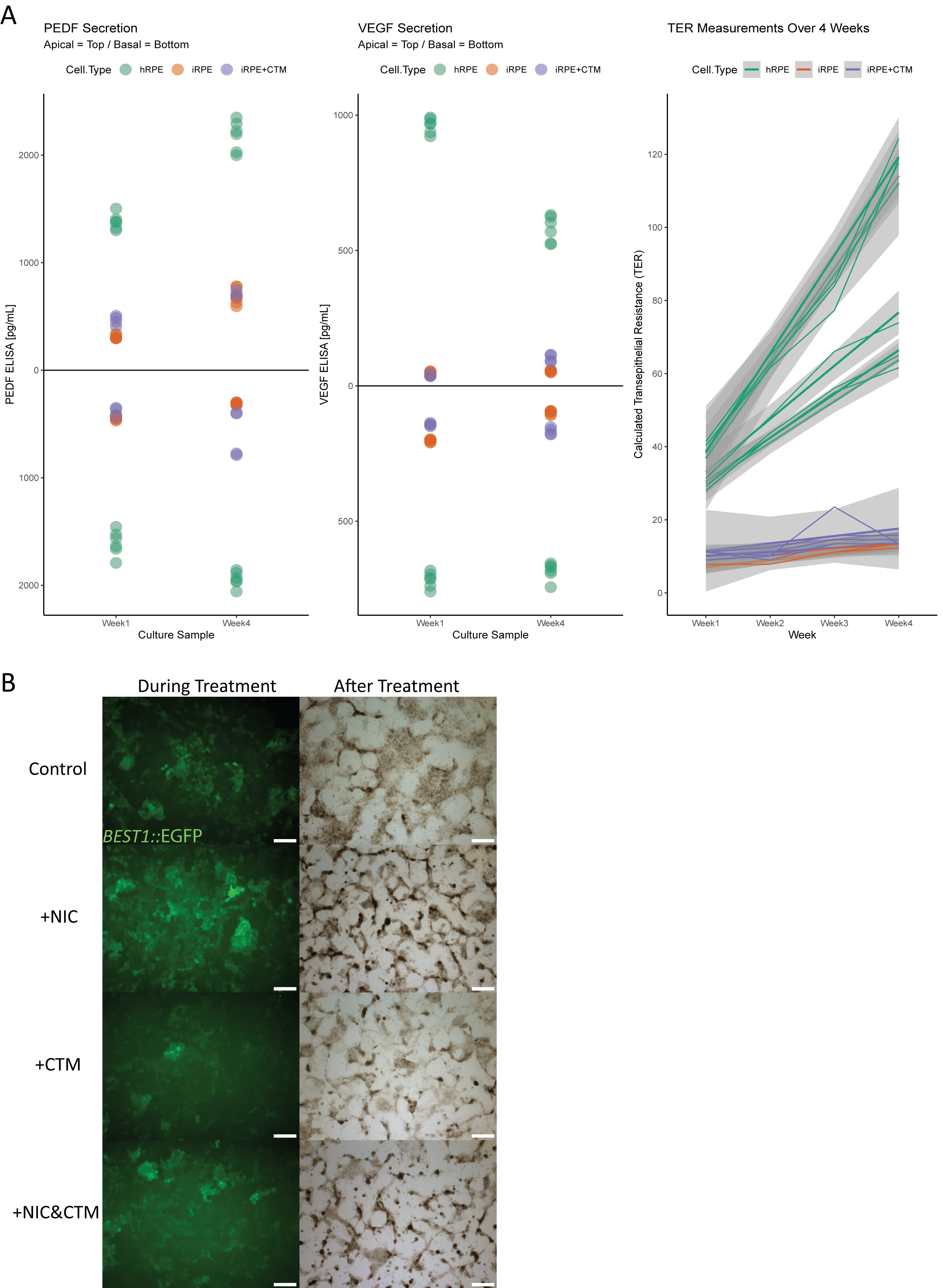

### Figure S4.png

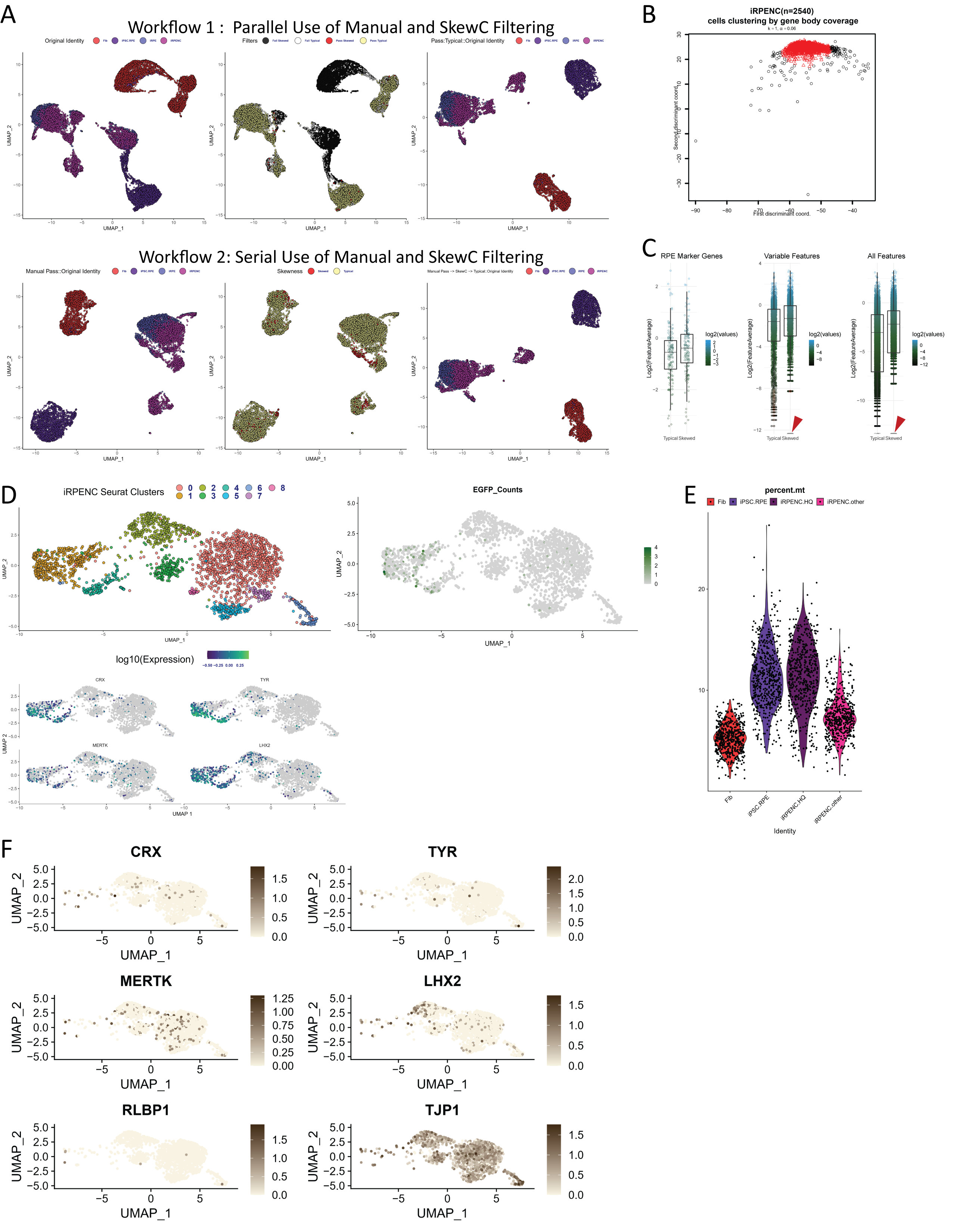

### Figure S5.png

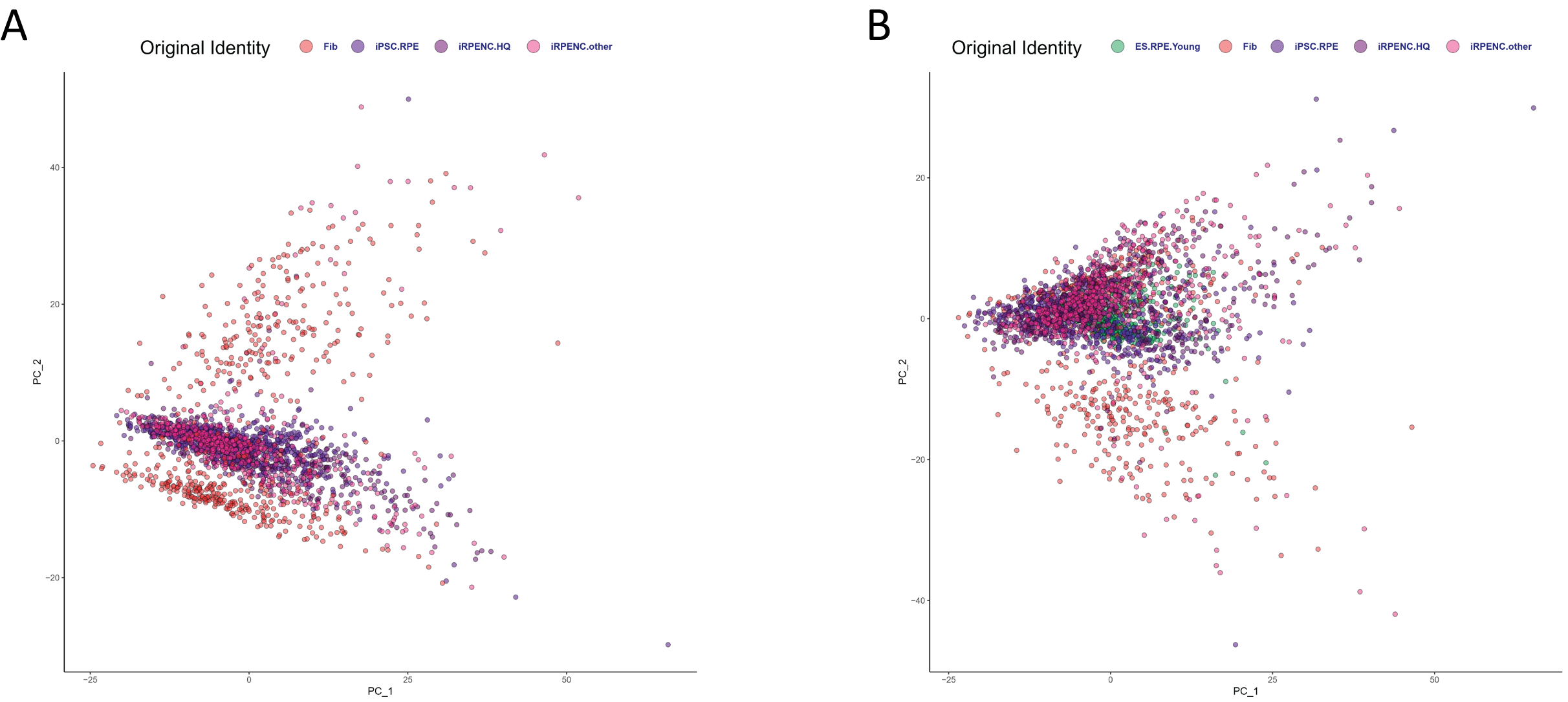

### Figure S6.png

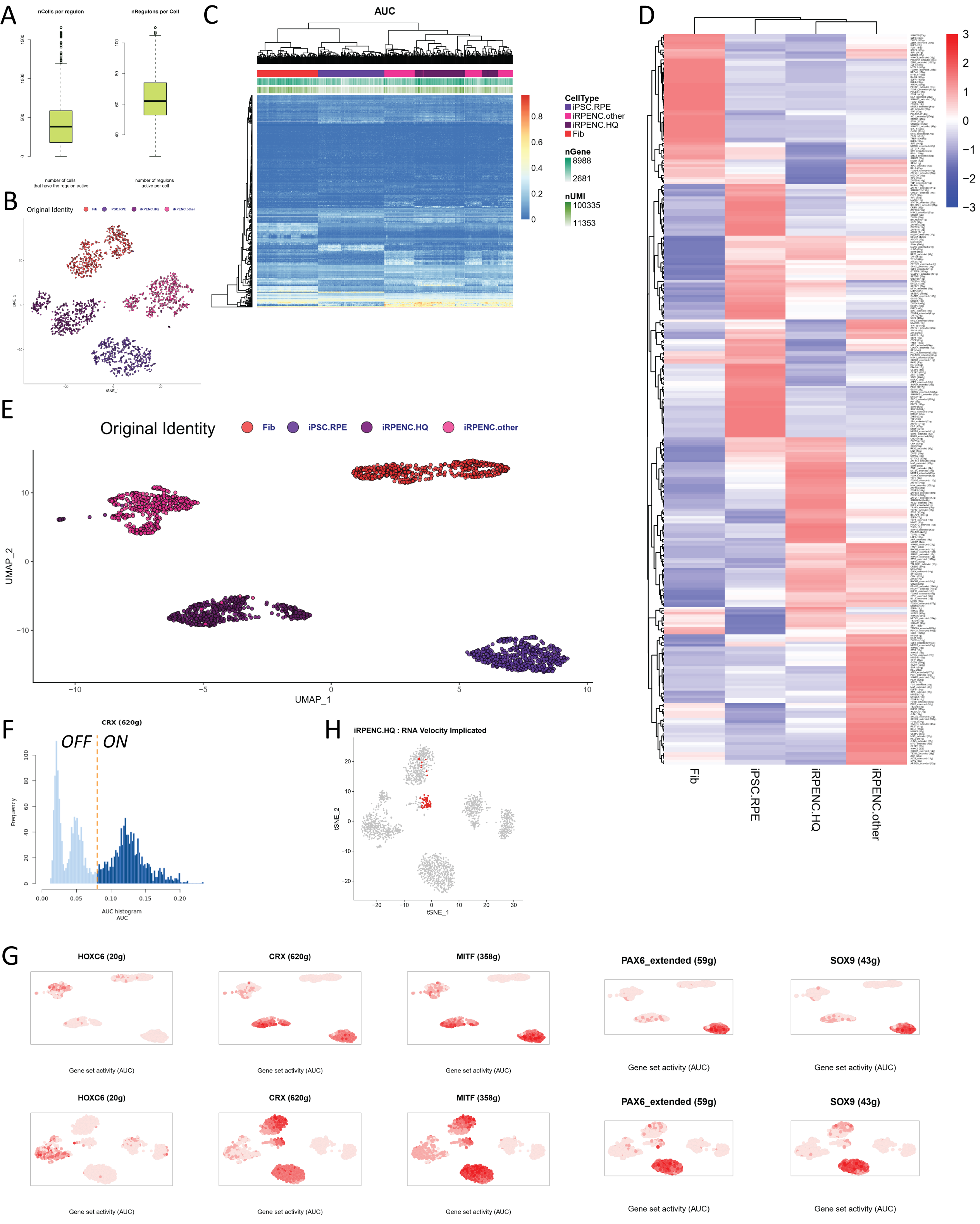
